# Nutritional effects on the expression of cryptic pigmentation in freshwater isopods

**DOI:** 10.1101/2025.05.13.653594

**Authors:** Moritz D. Lürig, Blake Matthews, Carsten J. Schubert, Osamu Kishida

## Abstract

Cryptic pigmentation is a key phenotypic adaptation that helps many invertebrates evade visual predators. However, little is known about whether and how the expression of pigmentation phenotypes that match the habitat background is influenced by the availability of nutritional resources. Here we investigated whether variation in both the background and the nutritive composition of benthic substrates affect the expressed pigmentation of freshwater isopods (*Asellus hilgendorfii)*. We collected isopods and their predominant substrate from 17 locations across Hokkaido, northern Japan, and quantified substrate background and nutritional composition (total protein, 18 amino acids, and C:N and C:P ratios). We found that variation in isopod pigmentation was better explained by the substrate’s nutritive composition than by its background darkness. Specifically, isopods were more pigmented when substrates had intermediate C:P ratios, lower C:N ratios, and a higher proportion of tryptophan - an essential amino acid involved in the isopods’ pigmentation pathway. These results are consistent with previous experiments showing that isopods reared under diets with higher protein concentrations developed more pigmentation, advancing our understanding about the environmental sources of phenotypic variation in natural populations. By demonstrating that nutritional constraints may shape the expression of key phenotypic adaptations in natural populations, our study opens new directions for exploring how organisms navigate adaptive landscapes; particularly in those organisms that rely on pigmentation for signaling and camouflage. Finally, we demonstrate how macronutrients, amino acids, and elemental ratios can serve as biotracers in ecological studies of adaptation, offering new opportunities to examine how stoichiometric traits influence phenotypic plasticity and adaptive capacity, especially in detritivorous taxa.

## Introduction

Pigmentation is a central phenotypic trait in most animals, serving crucial functions such as signaling, thermoregulation, and camouflage (Andrade & Carneiro, 2021; True, 2003). Especially for invertebrates with reduced motility or epibiotic habitus, such as benthic herbivores or detritivores, a close matching of cuticular pigmentation and background darkness is an important mechanism to avoid visual predation (Endler, 1978, 1992). Although behavior can influence habitat choice (Tavares et al., 2018) and thereby determine the value of a habitat as shelter (Lürig et al., 2016), there is only limited evidence for a coupling of behavior and pigmentation (e.g. through pleiotropy (Takahashi, 2013)). Instead, pigmentation in populations of detritivores is thought to be mostly shaped by adaptive evolution and visual predation (Clusella-Trullas & Nielsen, 2020). For many animals the genetic basis of pigmentation is well studied (Orteu & Jiggins, 2020), and there is increasing evidence for high heritability in pigmentation-related traits (Clusella-Trullas & Nielsen, 2020; Prokkola et al., 2013; Sheehan et al., 2017). Thus, under strong selection from visual predation, cryptic pigmentation may evolve rapidly to match a specific background darkness (Eroukhmanoff et al., 2009; Hargeby et al., 2004). However, the expression of pigmentation is known to be contingent on myriad environmental factors (Rautio & Korhola, 2002; Tollrian & Heibl, 2004), including resource availability (Britton & Davidowitz, 2022; Lürig & Matthews, 2021). Nevertheless, empirical studies examining this relationship in natural populations remain scarce, and our understanding of how environmental gradients in resource availability shape standing variation in pigmentation remains limited.

Ontogenetic development towards an organismal phenotype that matches the background can be dependent on resource availability particularly if adequate nutrition or precursor molecules are needed to generate pigmentation (Britton & Davidowitz, 2022). For example, genotypes with different developmental reaction norms may converge and diverge in their pigmentation at low versus high pigment availability, respectively. As such, pigment availability in a given environment might limit the range and nature of variation upon which predator-mediate selection can act. The evolutionary consequences of resource dependent variation in pigmentation also depend on the covariation between the coloration of the substrate background and pigment availability. Especially in situations where the ambient background is dark but resources to generate pigments are limiting, pigmentation could vary more strongly along gradients of nutrient availability in the substrate rather than its coloration. One common way to measure resource limitation of the substrate is by using essential elements (carbon, nitrogen and phosphorus) as a biotracer, which can vary strongly among substrates within and across ecosystems (Elser et al., 2000). Mismatches between consumers and their diets are common (Frainer et al., 2016; Halvorson et al., 2017; Martinson et al., 2008), and if they occur early in development they might be an important driver of phenotypic variation in invertebrates (Lürig & Matthews, 2021). Another way is to measure the external compounds that organisms cannot synthesize, such as essential amino acids (AA) or pigments themselves, such as carotenoids. At large, the physiological machinery to express pigmentation may be strongly genetically based (Clusella-Trullas & Nielsen, 2020; Prokkola et al., 2013; Sheehan et al., 2017), but a lack of adequate nutrition or essential compounds may impair development of pigmentation during early ontogeny (Lafuente & Beldade, 2019; Lürig & Matthews, 2021).

Freshwater isopods of the genus *Asellus* are members of the benthic detritivore community in freshwater ecosystems around the globe. These generalist detritivores can occur in very different types of habitats (creeks, ponds, lakes) where they feed on a large variety of benthic food items, such as allochthonous and autochthonous plant detritus, carcasses of con- and heterospecifics, or microbial colonies and fungal mats. This broad dietary spectrum often results in very high densities (up to 10000 animals per m^2^), making them an integral part of food webs in many freshwater ecosystems. Previous work on *Asellus aquaticus* has shown that there is considerable variation in the relationship of body size and pigmentation within and between populations across Europe (Eroukhmanoff et al., 2009; Hargeby et al., 2004; Lürig et al., 2019). Within populations, older isopods are larger, and larger isopods are more pigmented. Between populations, the strength of that relationship, i.e., the slope of pigmentation as a function of body size, can vary substantially. Two main drivers for this phenomenon have been considered: on the one hand there could be selection from visual predation for background matching pigmentation, i.e. crypsis (Hargeby et al., 2004, 2005), and on the other hand, there is evidence for diet-mediated developmental plasticity, namely increased pigmentation following ingestion of substrates containing the AAs required for pigment biosynthesis (Lürig et al., 2019; Lürig & Matthews, 2021). It seems plausible that both processes are closely interlinked through evolutionary and developmental processes, but while several field studies have investigated the crypsis hypothesis, none of these have tested the role of diet-mediated plasticity of pigmentation expression.

Populations of freshwater isopods in Japan represent a particularly promising opportunity to test these hypotheses, because i) many parts of the country are characterized by large networks of agricultural drainage channels and wetlands, exposing isopod populations to a wide array of nutritional conditions, and ii) many aquatic habitats in Japan experience strong predation pressure from invertebrate and vertebrate predators, such as dragonfly larvae, frogs, and salamanders (Kishida et al., 2009; Takatsu et al., 2016; Yamaguchi & Kishida, 2016). Using this comparative setup, we investigated the relationship between pigmentation in populations of freshwater isopods (*Asellus hilgendorfii*) across a range of different habitat and substrate types, and tested to what extent ambient gradients of nutrition and background darkness shape isopod phenotypes. We first conducted a field survey on Hokkaido Island, Japan, where we collected isopods and their associated benthic substrates from a range of different habitat types. We then quantified isopod pigmentation and background darkness using automated image analysis, and nutritional value of the habitat using chemical analyses, to investigate their relative contribution to variation in isopod pigmentation. Our results provide support for the hypothesis that the benthic substrates explain more the variation in isopod pigmentation in these natural populations than does substrate coloration. More generally, our findings highlight how the nutritive composition of resources can shape traits associated with crypsis, suggesting that variation in elemental ratios and amino acid concentrations may greatly influence how cryptic pigmentation is expressed in benthic invertebrates, and potentially, how it evolves.

## Methods

### Study system

Isopods of the genus *Asellus*, commonly referred to as “waterlouse” or “sowbugs”, are found in freshwater ecosystems throughout the northern hemisphere (Lafuente et al., 2021). *Asellus hilgendorfii* (Bovallius, 1886) is a freshwater crustacean that is part of a species complex of asellota isopods in eastern Asia (Sidorov & Prevorcnik, 2016), spanning parts of East Russia, Japan, and, since the 1970s, also in the western parts of North America (Toft et al., 2002). The species is morphologically similar to *A. aquaticus* (Linnaeus, 1758) found in Europe and Western Asia (Fig. 2), and both species are believed to form a complex of *s*pecies (*Asellus aquaticus-hilgendorfii)* across the palearctic realm (Sidorov & Prevorcnik, 2016). Throughout Japan *A. hilgendorfii* inhabits environments similar to those occupied by *A. aquaticus* in Europe (Miyashita & Yasuno, 1984), such as ponds, creeks, ditches and puddles with varying abundance of organic material (Hargeby et al., 2004; Lafuente et al., 2021). As such, *A. hilgendorfii* is considered an indicator of eutrophication in water bodies (Miyashita & Yasuno, 1984), and occupies a similar niche as *A. aquaticus* in Europe.

On Hokkaido island in Northern Japan, we observed that *A. hilgendorfii* exhibited similar behavior as *A. aquaticus*, i.e., individuals were sitting on detritus and consumed the substrate or the biofilm growing on top. This behavior likely makes them a common prey item to visual predators, such as salamanders (Takagi & Miyashita, 2019). Given the putative importance of cryptic pigmentation for survival, similar to *A. aquaticus, A. hilgendorfii* is an ideal study system to test the role of substrate background darkness versus its nutritional composition. To do so we collected samples of isopods and the substrates they were found on at 16 different sites across the island with varying environmental characteristics (Fig. 1A, Table 1). We quantified the relationship between body size and pigmentation as primary phenotypic variables, whose variation is visible with the bare eye. Additionally, we collected samples of benthic substrates representative of each site. In the laboratory, we quantified isopod pigmentation and benthic substrate darkness via imaging, and assessed the nutritional value of the substrates through chemical analysis.

**Table 1.** Overview of sampling sites. During our sampling campaign we collected a total of 1315 isopods, of which 1243 were large enough to be phenotyped using our computer vision pipeline (>2.5 mm). From each site we also took a representative sample of benthic substrate in exactly the same position where we collected the isopods. In the laboratory we used a combination of chemical analyses to quantify the concentration of 18 amino acids (=total protein, see Table S1 for details) and elemental ratios.

| Site number | Site name | Habitat type | Flow regime | Condition | N isopods | Background | Tryptophan (prop. t. protein) | Total protein (µg/g) | C:P ratio | C:N ratio |
| --- | --- | --- | --- | --- | --- | --- | --- | --- | --- | --- |
| 1 | Asahikawa 03 | Forest runlet | fast | pristine | 93 | 0.802 | 0.0022 | 0.059 | 342.401 | 23.506 |
| 2 | Atsushinai River | Forest runlet | static | pristine | 128 | 0.805 | 0.0004 | 0.043 | 498.864 | 42.530 |
| 3 | Kamekawa River | Forest runlet | static | pristine | 120 | 0.827 | 0.0270 | 0.035 | 381.584 | 23.706 |
| 4 | Asahikawa 02 | Pasture runlet | static | pristine | 14 | 0.824 | 0.0027 | 0.042 | 249.593 | 22.351 |
| 5 | Asahikawa 07 | Pasture runlet | slow | pristine | 17 | 0.804 | 0.0002 | 0.038 | 321.891 | 20.019 |
| 6 | Asahikawa 09 | Pasture runlet | fast | pristine | 121 | 0.808 | 0.0091 | 0.021 | 240.590 | 40.419 |
| 7 | Hiraoka Park | Pasture runlet | slow | pristine | 116 | 0.830 | 0.0003 | 0.033 | 164.285 | 39.204 |
| 8 | Teshio 03 | Pasture runlet | static | agriculture | 17 | 0.825 | 0.0003 | 0.026 | 289.328 | 29.055 |
| 9 | Asahikawa 01 | Roadside ditch | static | pristine | 62 | 0.688 | 0.0742 | 0.017 | 343.762 | 41.755 |
| 10 | Asahikawa 04 | Roadside ditch | static | pristine | 30 | 0.718 | 0.0186 | 0.038 | 361.969 | 55.568 |
| 11 | Nakahoronuka River | Roadside ditch | slow | agriculture | 73 | 0.714 | 0.0004 | 0.041 | 391.386 | 47.972 |
| 12 | Asahikawa 05 | Concrete channel | slow | agriculture | 132 | 0.764 | 0.0007 | 0.079 | 55.950 | 12.848 |
| 13 | Asahikawa 06 | Concrete channel | fast | agriculture | 73 | 0.802 | 0.0001 | 0.050 | 21.529 | 9.845 |
| 14 | Asahikawa 08 | Concrete channel | slow | agriculture | 92 | 0.757 | 0.0003 | 0.038 | 77.613 | 19.294 |
| 15 | Teshio 01 | Concrete channel | slow | agriculture | 99 | 0.807 | 0.0016 | 0.065 | 22.506 | 10.918 |
| 16 | Teshio 02 | Concrete channel | slow | agriculture | 56 | 0.767 | 0.0016 | 0.036 | 144.659 | 18.037 |

**Table 2.** Results for Linear Models. One linear model (LM) tested the relationship of isopod body size and pigmentation, and three linear mixed effect models (LMMs) tested for interactive effects of body size (logarithmized) and either habitat type (forest runlet, pasture runlet, roadside ditch, concrete channel), water flow regime (fast, slow, static) and environmental condition (pristine, agricultural), and site number as random effect. For LM 1, the first R2 value is the multiple and the second one the adjusted. For the LMMs, the first R2 is the conditional, and the second one the marginal.

| Model | Terms | df | F value / X<sup>2</sup> (RE) | P-value | R<sup>2</sup> (mult., adj.) |
| --- | --- | --- | --- | --- | --- |
| LM 1 | log(Length) | 1 | 1047.768 | <0.001 | 0.458, 0.457 |
| LMM 1 | log(Length) | 1, 1229.106 | 755.244 | <0.001 | 0.809, 0.275 |
|  | Habitat type | 3, 17.299 | 2.891 | 0.065 |  |
|  | log(Length) x habitat type | 3, 1228.835 | 14.469 | <0.001 |  |
|  | Site (RE) | 1 | 823.390 | <0.001 |  |
| LMM 2 | log(Length) | 1, 1230.285 | 909.215 | <0.001 | 0.806, 0.244 |
|  | Flow regime | 2, 17.767 | 4.110 | 0.034 |  |
|  | log(Length) x flow regime | 2, 1230.119 | 24.636 | <0.001 |  |
|  | Site (RE) | 1 | 848.728 | <0.001 |  |
| LMM 3 | log(Length) | 1, 1230.987 | 873.151 | <0.001 | 0.792, 0.253 |
|  | Env. condition | 1, 19.767 | 1.454 | 0.242 |  |
|  | log(Length) x env. condition | 1, 1230.987 | 22.675 | <0.001 |  |
|  | Site (RE) | 1 | 863.660 | <0.001 |  |

**Figure 1.**
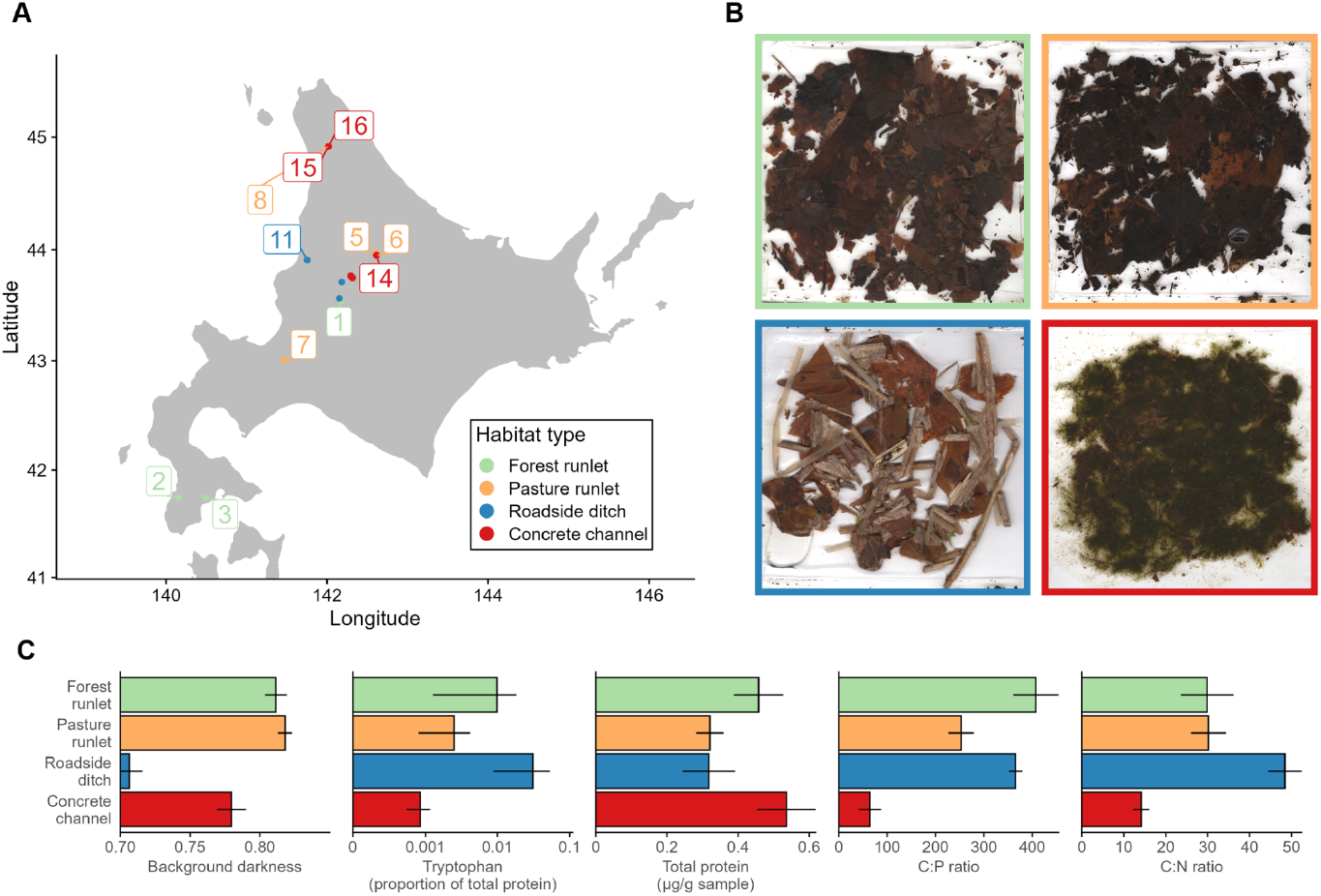
Overview of the sampling sites. **A:** The map shows our 16 sampling sites across Hokkaido, Japan, categorized into four types of habitat: forest runlets, pasture runlets, roadside ditches, and concrete channels. **B:** Exemplary samples of substrates for each habitat type that we obtained from the sites. We quantified brightness from the substrates using a modified flatbed scanner (Fig. S1), and nutritional profiles using a series of chemical analyses (Table 1). **C:** Average (mean ± standard deviation) background darkness, tryptophan concentration (proportion of total protein), total protein (per sample dry mass), ratios between carbon and phosphorus (C:P), and carbon and nitrogen (C:N).

### Isopod collection and phenotyping

We collected isopods between October 10th and 30th 2019 from the 16 sites (Table 1). We used dip nets to collect individual isopods directly from the benthic substrate. Immediately after collection we transferred the live isopods to aerated water tanks in the car. Within 48 hours of capture, we transported them to the laboratory and euthanized them by freezing at -20°C. We then thawed them again 1-2 days later to take high resolution images using a modified flatbed scanner (Epson V37), following the protocol of Lürig et al. (2019). Specifically, we placed isopods dorsally on the slightly moistened surface inside the scanner (Fig. S1), and scanned all isopods within a single site at 600 DPI. We then analyzed the images with computer vision: using the Python package phenopype (Lürig, 2022), we segmented the images into foreground (individual isopod specimens) and background. From each detected isopod (excluding legs and antennae), we extracted body size as the diameter of the minimum enclosing circle, (i.e., excluding legs and antennae), and mean pixel intensity after converting from three-channel RGB to single-channel grayscale images. We then converted the extracted grayscale values (0=black, 255=white) to an inverted 0-1 scale (0=white, 1=black).

### Analyses of habitat substrates

To quantify the effects of background darkness and nutritional quality on isopod size and pigmentation we also took samples of the benthic substrate from which we collected isopods. We took three samples per site that were representative of the isopod microhabitat by either collecting or scraping the substrate with the dip nets and storing it in plastic bags in darkness and cooled. After transporting the samples to the laboratory, we quantified the darkness of the substrate in the same fashion as we did for isopods, i.e., using the same modified flatbed scanner and by analyzing the resulting images with computer vision. Specifically, we separated a portion from the bulk and spread it out on the wetted scanner surface, segmented the substrate from the scanned images (i.e., excluded all white background), and quantified mean background darkness on the same 0-1 scale as isopod pigmentation (0=white, 1=black).

When analyzing the nutritional quality of the substrate we considered two aspects: the stoichiometric ratio of major elements (C, N, P) and the concentration of 18 AA (see Table S1 for details). Elemental ratios provide information on quality and value of substrates as diet (Elser et al., 2000; González et al., 2017; Jeyasingh et al., 2014; Leal et al., 2017), whereas AAs provide a more nuanced view of protein content and individual AAs that might be essential, i.e., cannot be synthesized by the organism (Pickett et al., 2024). In our analyses we focussed the proportion of tryptophan among all measured AA (hereafter tryptophan concentration), the initial precursor molecule in the synthesis of the main integumental pigment in isopods (xanthommatin)(Needham & Brunet, 1957; Oetinger & Nickol, 1982), and on the summed concentration of all 18 AAs (hereafter total protein). To measure the concentration of C and N we used an elemental analyzer, while we used a continuous flow analyzer to measure P concentration (see SI for details). We scaled element concentrations by the molar mass of the respective element prior to calculating elemental ratios. To measure the concentration of AAs we followed the protocol from Saboret et al. (2023) (see SI for details).

### Statistical analyses

We conducted all statistical analysis and made all plots with the statistical programming language R version 4.3.3 (R Core Team, 2024). First, to test the relationship of size and pigmentation with body size (logarithmized - also in all other analyses) we used a linear model (LM 1, Fig. 2B). With the residuals of LM 1 we visually inspected within and among site and habitat variation in size-corrected pigmentation (Fig. 2D). Second, to test whether and how discrete environmental descriptors would predict the relationship between body size and pigmentation, we used a series of three linear mixed effect models with interaction terms of body size and habitat type (LMM 1, Fig. 2C), water flow regime (LMM 2, Fig S2A) and environmental condition (LMM 3, Fig S2B), and with site number as random effect. We used the predictions of LMM 1 to assess variation in pigmentation among habitat types (Fig. 2E). We implemented the LMMs using the *lme4* package v(1.1-35.1) (Bates et al., 2015).

**Figure 2.**
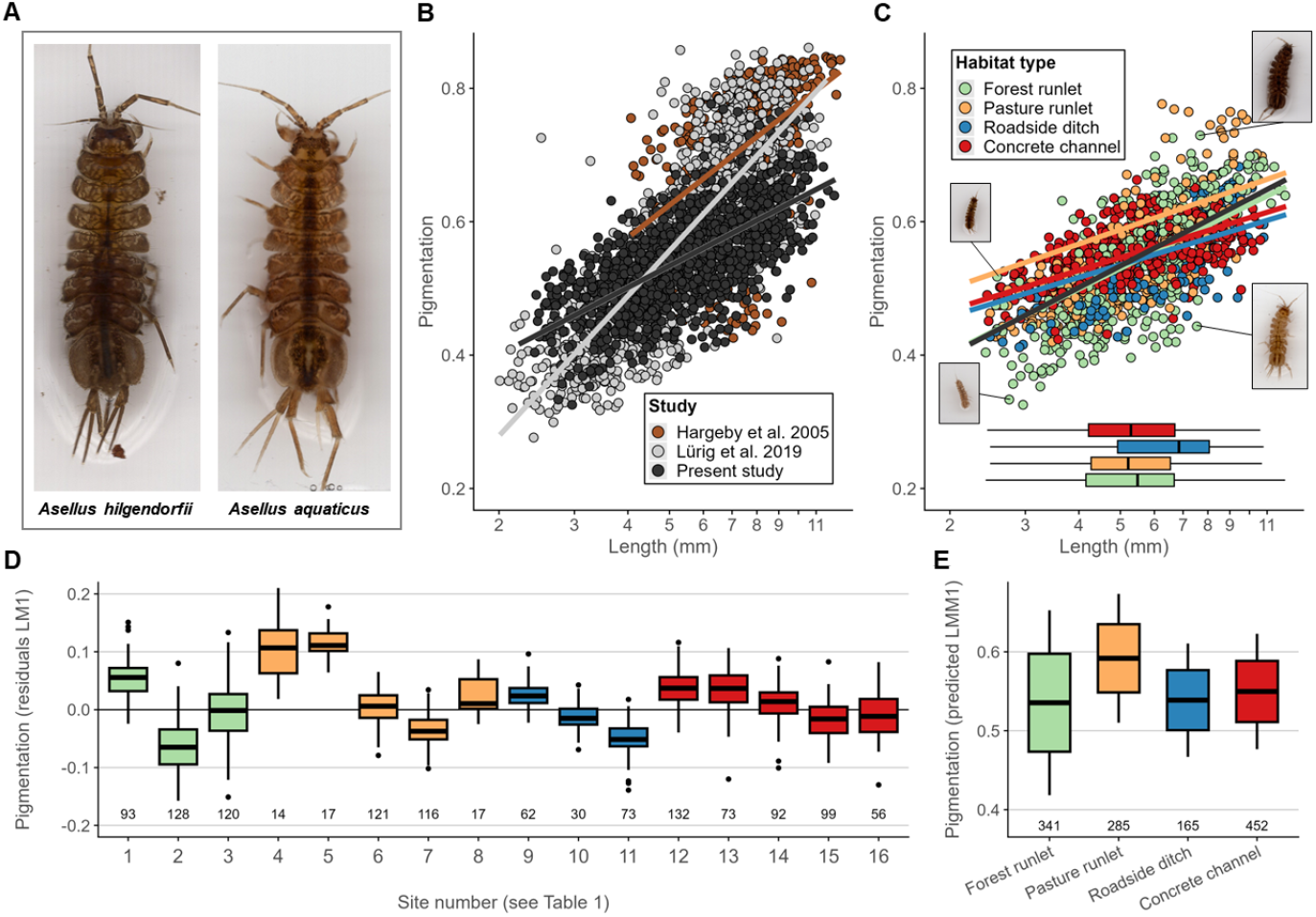
The relationship of body size and pigmentation in isopods. **A:** *Asellus hilgendorfii*, collected on Hokkaido island, Japan, and *Asellus aquaticus*, collected from Lake Lucerne, Switzerland. **B:** The relationship between body size (log transformed) and pigmentation in *A. aquaticus* from previous studies (Hargeby et al., 2005; Lürig et al., 2019) and *A. hilgendorfii* from this study. As isopods grow, they develop more cuticular pigmentation, so that the largest isopods are typically also the most pigmented ones. Thus, pigmentation in Asellidae has to be considered together with body size (e.g., by using a regression to extract size-corrected residuals, or by looking at slope and intercept). The individuals of *A. hilgendorfii* we collected on Hokkaido (N=1243) have a more shallow slope than Swiss and Swedish *A. aquaticus*, but are overall similar in their relationship of size and pigmentation (black line: fit from linear regression [LM 1, Table 2]). **C:** On Hokkaido, the relationship between size (log transformed) and pigmentation is almost identical in all habitats, except agricultural concrete channels, where the slope is more shallow but the intercept is bigger. (colored lines: fit from linear regression [LMM 1, Table2], black line: fit from LM 1). Horizontal boxplots at the bottom show the distribution of body sizes per habitat type. **D:** Boxplots of residuals from LM 1 (black line in panel B and C), grouped by habitat type; numbers below are N isopods per site (see Table 1). **E:** Model estimates for pigmentation by habitat (LMM 1). There is a significant interaction between body size and habitat type, which, however, only explains a small amount of variation in pigmentation we observed across the different sites, and within the habitat types (panel D).

Third, to test whether and how continuous biotic and abiotic environmental gradients would modify the relationship of body size and pigmentation, we used a series of five Generalized Additive Models (GAM). We chose to use GAMs rather than LMMs because we expected complex relationships between environment and phenotypes, e.g., elemental ratios (Leal et al., 2017). Each GAM used thin plate spline smooth terms to test body size and a single covariate: either background darkness or one of four aspects of substrate nutritional value (tryptophan concentration, total protein, C:N ratio and C:P ratio) on isopod pigmentation. In addition, each model used a single tensor product interaction term with body size and the covariate. All models used sites as a random effect and five knots for the smoothing. We implemented the GAMs using the *mgcv* package (v1.9-1) (Wood, 2011). It is important to note that while the GAM framework offers a frequentist approach to testing significant contributions of smoothing terms to the explained variation in the response variable, the resulting F-values reflect the reduction in unexplained deviance and do not provide a direct measure of effect size or relative importance, similar to their interpretation in linear models. Therefore, visual interpretations are necessary to assess the relative differences in strengths between the variables. We therefore visualized the model predictions for pigmentation in multivariate space along axes of body size and the different covariates (Fig. 3A), as well as univariate predictions for pigmentation over a range of body size for 10th, 50th, and 90th percentiles of the five covariates (Fig. 3B). Finally, to summarize and compare the relative importance of the different covariates in explaining pigmentation, we performed a hierarchical partitioning with all variables (Fig. 4).

**Figure 3.**
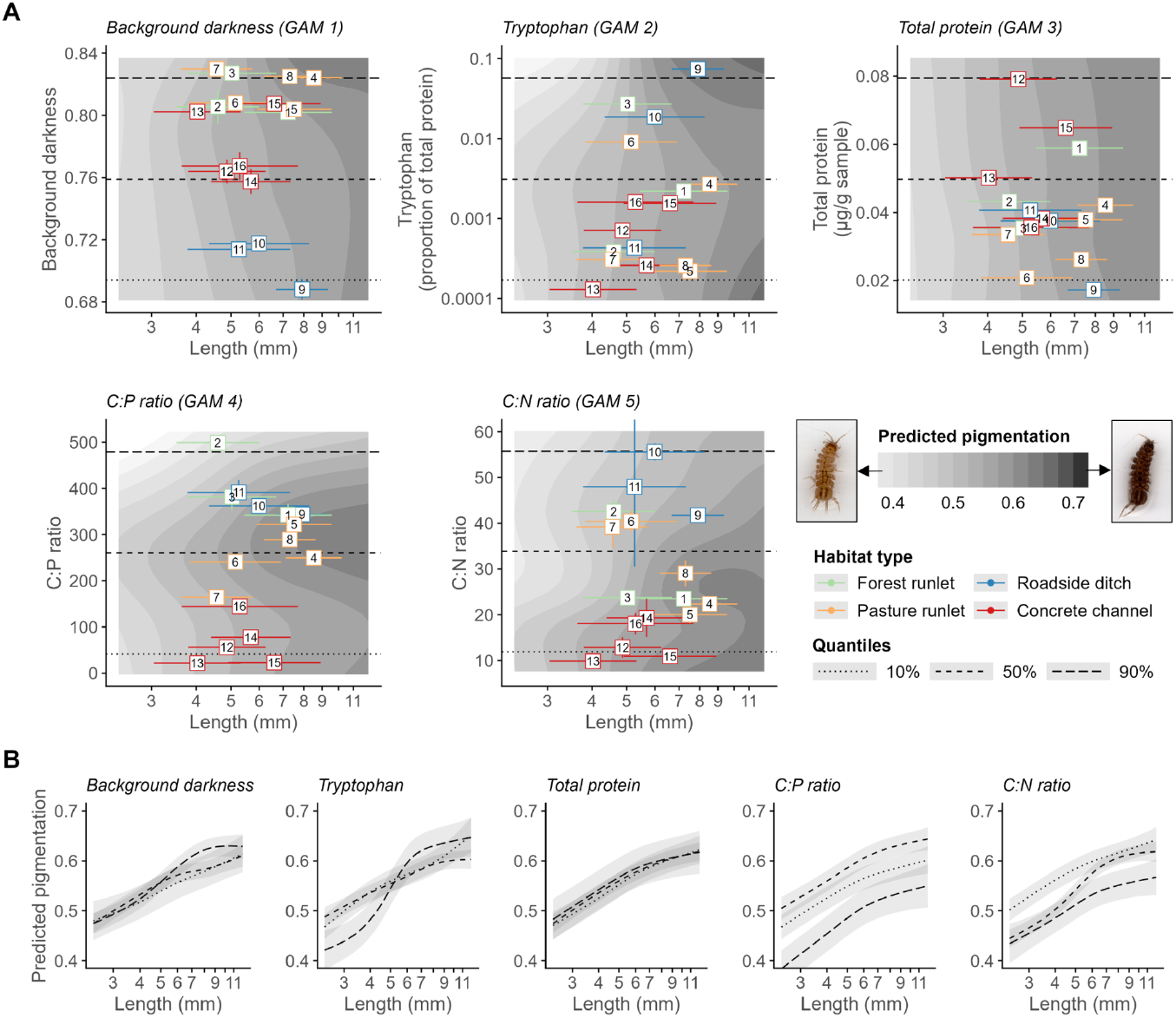
The relationship of body size and pigmentation is affected by background and nutrition. A: We used a series of Generalized Additive Models (GAM) to test whether the relationship of body size and pigmentation is modified by background darkness or aspects of nutrition. Each model used main and interactive smooth terms of body size and either background darkness (GAM 1), tryptophan concentration (GAM 2), total protein (GAM 3), C:P ratio (GAM 4) or C:N ratio (GAM 5). The gray-shaded surfaces in each panel are estimates for pigmentation from the interaction term of each model. The horizontal error bars are mean and standard deviation of body size for each of the 16 populations (indicated site numbers listed in Table 1), the vertical error bars are mean and standard deviation of the technical replicates in background darkness, C:N and C:P (N=3; we had only one technical replicate for amino acids). Across all variables, pigmentation generally increases with body size (i.e., darker shades of gray from left to right along the x-axis), but especially variation in C:P and C:N ratio can strongly influence this relationships: with intermediate to low elemental ratios, isopods become darker for a given body size. For details on model fits see Table 3. **B:** Predicted pigmentation (mean ± standard error) for fixed values (10th, 50th, and 90th percentile, dashes correspond to the vertical lines in panel A) of the covariates from the GAMs. Only the darkest backgrounds were associated with higher than expected pigmentation in large isopods. Total protein did not significantly affect the relationship of body size and pigmentation (all confidence intervals overlap). High tryptophan concentration results in a much steeper pigmentation-body size relationship. Intermediate C:P ratios result in a higher intercept than low or high ratios, whereas low C:N ratio results in higher intercepts.

**Figure 4.**
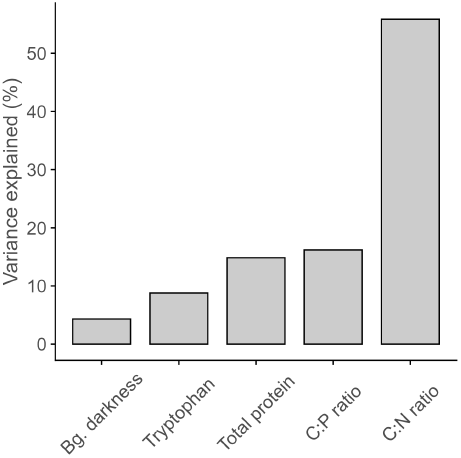
Variance partitioning of pigmentation. Hierarchical partitioning, which quantifies the independent contribution of each predictor to the explained variance, confirms the strong relative contribution of nutritional aspects of the benthic substrate and especially C:N ratios to variation in pigmentation in Japanese isopods. We used size corrected pigmentation for the test (residuals from LM 1)

## Results

Across all sampled sites, individuals of *A. hilgendorfii* collected from Hokkaido island exhibit variation in the relationship of body size and pigmentation to a similar extent as *A. aquaticus* from populations in Sweden and Switzerland (Fig. 2): larger individuals have stronger pigmentation than smaller individuals (Table 2, LM 1). The slope of this relationship is similar to populations sampled in southern Sweden reported in Hargeby et al. (2005) (Fig. 2B), but more shallow than the one from Swiss populations from Lake Lucerne reported in Lürig et al. (2019). We found substantial variation in size-corrected pigmentation within a given habitat type (Fig. 2D, residuals of LM 1), and also between the different habitat types (Fig. 2E, predictions of LMM 1). Specifically, isopods collected from pasture runlets have higher predicted pigmentation than all other habitats (i.e., highest intercept), whereas isopods from forest runlets have the biggest spread across all body sizes (i.e., steepest slope). However, both the substantial variation within a habitat type (Fig. 2D) and the large contribution of the random effects in the LMMs (Table 2) indicates the relationship of body size and pigmentation is neither well explained by habitat type, nor by water flow regime (static, slow flowing, fast flowing) or environmental condition (pristine, agricultural).

**Table 3.**
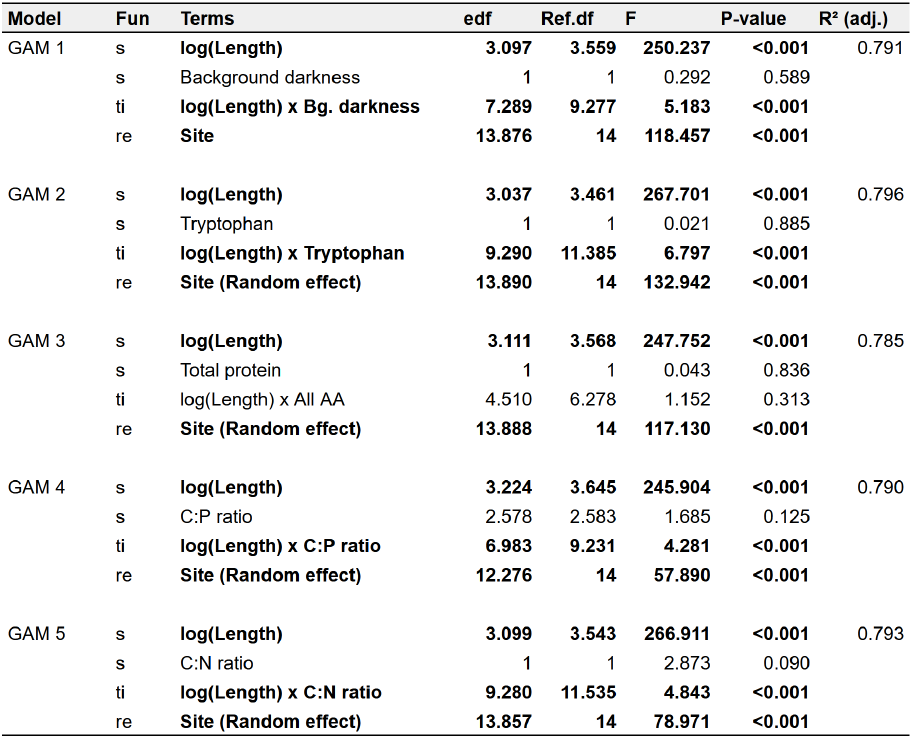
Results for Generalized Additive Models. The table shows the results for the five Generalized Additive Models (GAM). Fun indicates the smooth function used for the different terms (s = thin plate spline, ti = tensor product interaction, re = random effect). Edf are estimated and Ref.df reference degrees of freedom. Significant results (P-value > 0.05) are highlighted in bold.

All continuous biotic and abiotic environmental gradients except total protein affected the relationship of body size and pigmentation, as indicated by the significant interaction term in the GAMs (Table 1) and the multi- (Fig. 3A) and univariate visualizations (Fig. 3B). The elemental ratios had particularly strong effects: at intermediate C:P ratios, isopods were most pigmented for a given body size, where high or low ratios were associated with weaker pigmentation. Moreover, substrates with low C:N ratios had the highest intercept, whereas intermediate C:N ratios resulted in steeper slopes. Only a high tryptophan proportion in total protein was associated with stronger pigmentation for a given body size, indicated by the steeper slope than intermediate or low concentrations. Background darkness had only a minor effect: only at sites with very dark backgrounds, large isopods tended to be darker than similarly sized isopods in lighter backgrounds. Overall, C:P ratios explained most of the variation in size-corrected pigmentation relative to all other covariates we tested (Fig. 4).

## Discussion

Cryptic pigmentation is an important phenotypic adaptation of benthic invertebrates to avoid predation from visual predators (Hargeby et al., 2005; Lürig et al., 2019; Merilaita, 1998), but it remains poorly understood whether environmental mismatches in resource availability and quality constrain the expression of matching pigmentation phenotypes (Petrullo et al., 2023; Twining et al., 2022). Here, we examined how pigmentation in freshwater isopods responds to both background darkness and the nutritional composition of benthic substrates used as microhabitat and dietary resource. While pigmentation increased with darker backgrounds, dietary elemental ratios had a far stronger influence on body size–pigmentation relationships. Thereby our results confirm and expand findings from previous field surveys (e.g. Hargeby et al., 2005) and experimental work (e.g. Lürig & Matthews, 2021), and - for the first time - demonstrate variation in pigmentation in isopod populations can be partially explained by quantified variation in nutritionally-relevant chemical composition of available resources. The collective evidence from these multiple lines of research indicates that crypsis, while likely an important factor in shaping phenotypes of isopod populations via natural selection from visual predation, might be contingent on the presence of sufficient nutritional resources, here quantified as stoichiometric ratios of C,N and P, and the concentration of essential AAs.

Individuals of *A. hilgendorfii* become more pigmented with increasing body size (Fig. 2), reflecting the same ontogenetic process of pigment accumulation with age as in *A. aquaticus* (Lafuente et al., 2021; Lürig et al., 2019; Needham & Brunet, 1957). However, the relationship of body size and pigmentation in Japanese populations was not as steep as in populations of *A. aquaticus* from Sweden and Switzerland: especially the larger specimens of *A. hilgendorfii* are less pigmented for a given body size (Fig. 2B). One possible explanation is that habitats in Japan were not as dark as the habitats previously sampled in Europe. In Sweden, the most pigmented isopods were found in stands of reed (*Phragmites australis*) (Hargeby et al., 2005), which produces a surface layer of extremely dark detritus as it decays, while many of the darkest Swiss specimens were collected from ponds with black alder foliage (*Alnus glutinosa*) (Lürig et al., 2019), which is darker than most broad-leafed trees in Japan. The swedish studies also focussed on sampling from habitats with strong contrasts in background darkness, i.e., light-green macrophytes (*Chara tomentosa*) on bright sand vs. dark brown stands of reed on black detritus (Eroukhmanoff et al., 2009; Hargeby et al., 2004, 2005). In contrast, our approach was to sample a broader range of habitats with more gradual differences in background darkness. This may explain why we only found a relatively weak effect of both discrete habitat type (LMM 1) and continuous background darkness (GAM 1) on the relationship of body size and pigmentation. Although our findings differ from those of the Swedish work, which found strong effects of background darkness of different habitats on pigmentation, it does not exclude the importance of crypsis for survival. However, it indicates that other factors may be more important in determining the relationship of body size and pigmentation along spatial gradients.

The lack of effect of total protein concentration on the relationship of body size and pigmentation indicates that macronutrients like bulk dietary protein are not crucial for pigment synthesis in asellid isopods. This finding is somewhat surprising, because it does not fully align with the results from previous experimental work that found more pronounced development of pigmentation during ontogeny under high vs. low dietary protein supply (Lürig et al., 2019; Lürig & Matthews, 2021). However, the aforementioned experiments did not manipulate protein concentration independently of tryptophan, an essential precursor AA for pigment synthesis in isopods (Figon et al., 2020; Linzen, 1974). It is possible that the differences in pigmentation rate and strength observed in these experiments may have been due to differences in tryptophan, and not protein concentration. In contrast, the sites in our field survey varied independently in both tryptophan and total protein concentration (Fig. 1C): we sampled sites with high tryptophan but low protein concentration (e.g., site 9), and vice versa (e.g., site 12). While the sites with low protein and high tryptophan did not host isopods with higher than expected pigmentation, those with high protein and low tryptophan did. Only for substrates with very high proportions of tryptophan among total protein content (e.g., site 9 and 3) was there a significantly steeper slope than the mean for all populations combined (Fig. 3B). Importantly, the sites with the darkest isopods for a given body size (i.e., site 4 and 5) had relatively low tryptophan concentration. Unless the isopods also consumed substrates from adjacent habitats with higher tryptophan concentration, our finding suggests that already very low concentrations of tryptophan are sufficient for xanthommatin synthesis in isopods, highlighting the important but complex role of specific external dependencies in determining phenotypic variation (Badyaev et al., 2019; Pickett et al., 2024).

The ratios of major elements had a clearly discernible effect on the relationship of body size and pigmentation across all habitats and sites: intermediate C:P (250-350) and somewhat lower C:N ratios (20-30) were associated with the darkest pigmentation for a given body size. These numbers are in line with previous experimental research (Lürig & Matthews, 2021), and a large body of work suggesting that consumers tend to align the stoichiometric relationship of essential elements between themselves and the resources they consume (Elser et al., 2000; González et al., 2017; Jeyasingh et al., 2014; Leal et al., 2017). Indeed, the substrates that result in strong pigmentation for a given body size have elemental ratios that are more similar to isopods than other substrates associated with lighter pigmentation (Lürig & Matthews, 2021). Interestingly, the sites with intermediate elemental ratios and strongly pigmented individuals (i.e., 1, 4, 5, 8 and 9), also had only moderate levels of protein content. This could be explained by a previous finding, where high dietary protein resulted in higher mortality (Lürig & Matthews, 2021), putatively due to the costly nature and harmful byproducts of protein digestion (Arganda et al., 2017; Le Couteur et al., 2016; Linzen, 1974). Despite their undoubtedly strong effect on pigmentation, intermediate elemental ratios were not always associated with strong pigmentation. For instance, site 9 had very dark isopods, but also the lightest background and elemental ratios that are not optimal for strong pigmentation. However, site 9 also had the highest tryptophan concentration (by an order of magnitude), which may have led to the stronger than expected pigmentation. Overall, these findings suggest that elemental ratios, particularly C:N (Fig. 4), are good predictors of pigmentation phenotypes in the wild, but interactions with AAs may further modulate their expression, underscoring the complex relationship between dietary composition, microhabitat substrate darkness, and phenotypic traits in natural populations.

We are only beginning to understand how intraspecific variation in phenotypes is shaped by how organisms navigate their nutritional landscapes. Our findings are consistent with a growing body of work that uses biotracers to study ecological processes and evolutionary dynamics through the lens of nutritional phenotypes (Leal et al., 2017; Pickett et al., 2024). For instance, it is now known that spatio-temporal variation of biotracers such as elemental ratios (Elser et al., 2000; González et al., 2017; Leal et al., 2017), AAs (Dwyer et al., 2018, 2019) or micro- (e.g., fatty acids (Saboret et al., 2023; Twining et al., 2021)) and macronutrients (e.g., proteins and lipids (Le Couteur et al., 2016; Solon-Biet et al., 2015)) can affect consumer performance, and thereby play a role in structuring food webs (Thompson et al., 2015), regulating ecosystem function (Hillebrand et al., 2009), and determining community composition (Tumolo et al., 2023). However, only seldomly are the direct effects of biotracers on phenotypes and developmental processes been quantified in natural populations (but see Katayama et al., 2016). In this context, our study goes beyond the current state of the art by showing that elemental ratios and AA availability jointly influence pigmentation phenotypes among natural populations, and our results suggest there may be some nutritionally dependent constraints on the ability of organisms to express phenotypes associated with the presumably high adaptive value of cryptic pigmentation in a particular habitat. This suggests that nutritional dimensions of adaptive landscapes may warrant further investigation, particularly if we want to identify the underlying ecological causes of natural selection in the wild. Our work further highlights that biotracer variation, which has mostly been studied at the species level (González et al., 2018), is increasingly useful for studying intraspecific variation in stoichiometric and AA traits (Jeyasingh et al., 2014; Lemmen et al., 2019). We therefore recommend that future research investigates how intraspecific variation in stoichiometric traits shapes phenotypic plasticity and adaptive potential, particularly in emergent study systems like freshwater isopods (Lafuente et al., 2021), which serve as key components of food webs and are highly sensitive to variation in elemental ratios (Evans-White & Halvorson, 2017).

## Conclusions

Our study shows that the composition of major elements and AAs in nutritional resources rather than background darkness can better explain the expression of cryptic pigmentation in natural populations of freshwater isopods. These results indicate that the expression of adaptive phenotypes, i.e., pigmentation that matches background darkness of microhabitats with strong visual predation, might be constrained by nutritional requirements, such as optimal elemental compositions or essential This suggests that in addition to characterizing the adaptive landscape, we also need to consider the factors that allow organisms to navigate it - for instance, through experimental manipulations of biotracers such as elemental ratios, AAs, and micro- and macronutrients, which provide the resources for the expression of a particular phenotype during ontogeny. In light of this, dietary mismatches may hinder organisms in expressing phenotypes that increase their fitness in specific environments, which may have consequences for selective processes and evolutionary dynamics.

## Acknowledgments

We would like to express our sincere thanks to Kotaro Takai for leading the field campaign on Hokkaido and for sharing his in-depth knowledge of the local natural history and the ecology of isopods. Moreover, we thank Daniel Steiner for his expertise in the chemical analysis of substrate samples, and Grégoire Saboret for insightful conversations about amino acids.

## Author Contributions

MDL and OK conceived and designed the study. MDL carried out the field work, analyzed the data, and wrote a first draft of the manuscript. BM contributed to data analysis. CS organized and provided feedback on chemical analyses. All authors contributed substantially to interpretation of the results and writing of the manuscript, approved the final version, and agreed to be accountable for the integrity of their work.

## Statement on inclusion

Our study brought together researchers from Europe and Japan to investigate ecological processes in freshwater habitats on rural Hokkaido. This collaboration not only integrated region-specific ecological knowledge but also provides insights that may benefit local communities by enhancing the understanding of biodiversity and ecosystem dynamics in the region’s freshwater systems.

## Data availability statement

All data and code necessary to reproduce the results and figures shown in this paper are available via Zenodo (Lürig et al., 2025).

## Supplementary Information

### Methods

#### Chemical analysis of habitat substrates

Prior to the chemical measurements we thoroughly dried (drying oven at 60°C over 48 hrs) and homogenized the samples using mortar and pestle. To quantify C, N, and P content we filtered each benthic substrate sample onto two ashed (400°C), pre-weighed Whatman GF/F glass microfiber filters (47 mm), dried them overnight at 60°C, and reweighed them to determine total dry mass. We measured C and N content from one filter using an Elementar vario PYRO cube EA-IRMS, and measured P content from the other filter using a Skalar San++ Continuous Flow P/N analyzer, after digesting the samples with a peroxydisulfate solution and diluting them (1:20).

To quantify AAs we took a subsample from the homogenized sample that weighed between ∼100 and ∼250 mg (average = 198,9 ±12.03). We then dried the material at 60°C for at least two days and further homogenized it with tungsten beads (3 min at 30 Hz), and afterwards hydrolyzed the material with 1–2 mL 6 M HCl at 110°C for 20 h. We prepared an in-house standard for all 18 AAs using >99.5% pure individual AAs from Sigma-Aldrich and derivatized free AAs to N-acetyl methyl ester (NACME-AA) derivatives following Corr et al. (2007) and Larsen et al. (2013). We then extracted lipophilic compounds with 2 mL n-hexane/dichloromethane, added 0.1–1 mL (depending on sample concentration) of norleucine (Nle) 4.7 μMol as an internal standard, and evaporated the HCl from hydrolyzed samples at 110°C under N_2_ before derivatization. We then methyl-esterified carboxyl groups with acidified methanol (6:1 anhydrous methanol: acetyl chloride) for 60 min at 75°C, then acetylated amine groups with an acetylation mixture (1:2:5 acetic anhydride: triethylamine: acetone) for 10 min at 60°C. We flushed vials with N_2_ before and after reactions and dried excess solvent under N_2_ at 60°C. We dissolved samples in 2 mL ethyl acetate, removed precipitates and salts with 1 mL saturated NaCl solution, and dried the ethyl acetate solution under N_2_. We brought the final volume to ∼300 μL in gas chromatography (GC) vials. We separated amino acids on an Agilent 6890 Series GC System with a Gerstel MultiPurposeSampler MPS2 and an InertCap 35 GC column (GL Sciences, 60 m × 0.32 mm × 0.50 mm). We programmed the GC oven as follows: 50°C (start), 15°C/min to 140°C, 3°C/min to 152°C (held 4 min), 10°C/min to 245°C (held 10 min), and 5°C/min to 290°C (held 5 min).

## Figures

**Figure S1.**
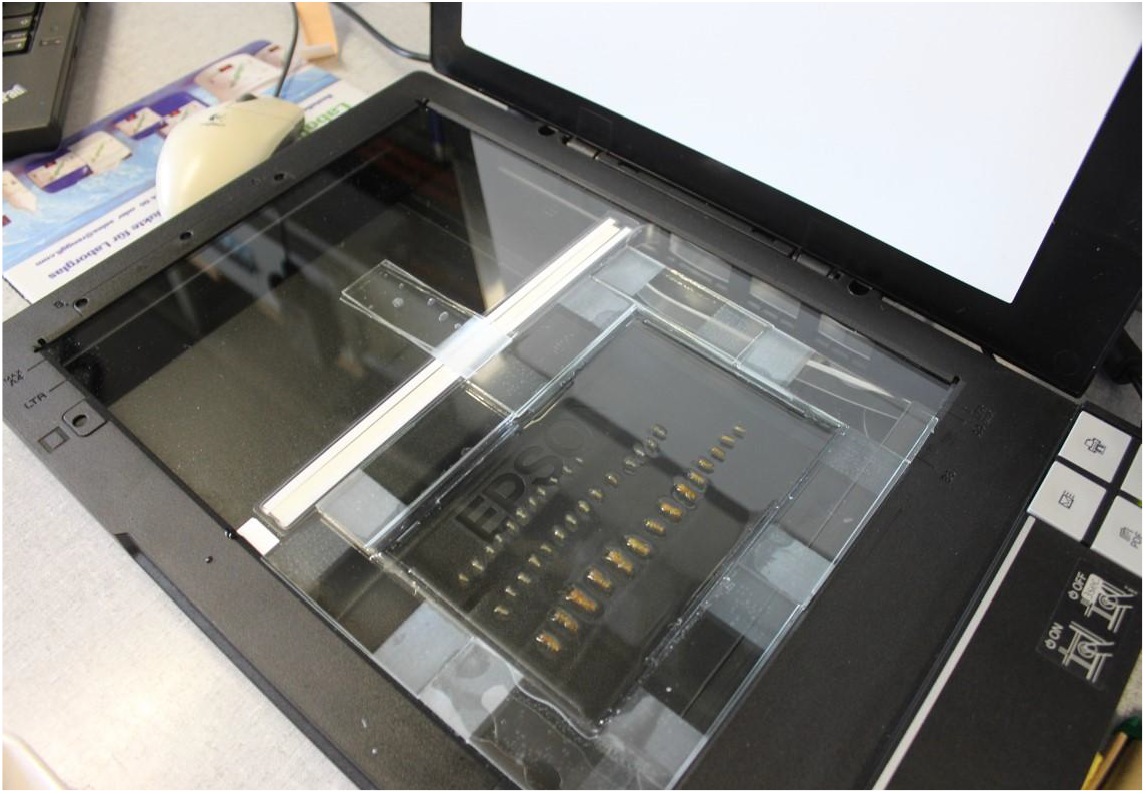
Flatbed scanner used to scan isopods. We used a modified document flatbed scanner (Epson V37) to scan isopods and habitat substrates. To scan the substrates and isopod specimens we covered the surface inside the glass-rectangle with water, and used a drop of soap to break the surface tension. For a detailed protocol refer to Lürig et al. (2019).

**Figure S2.**
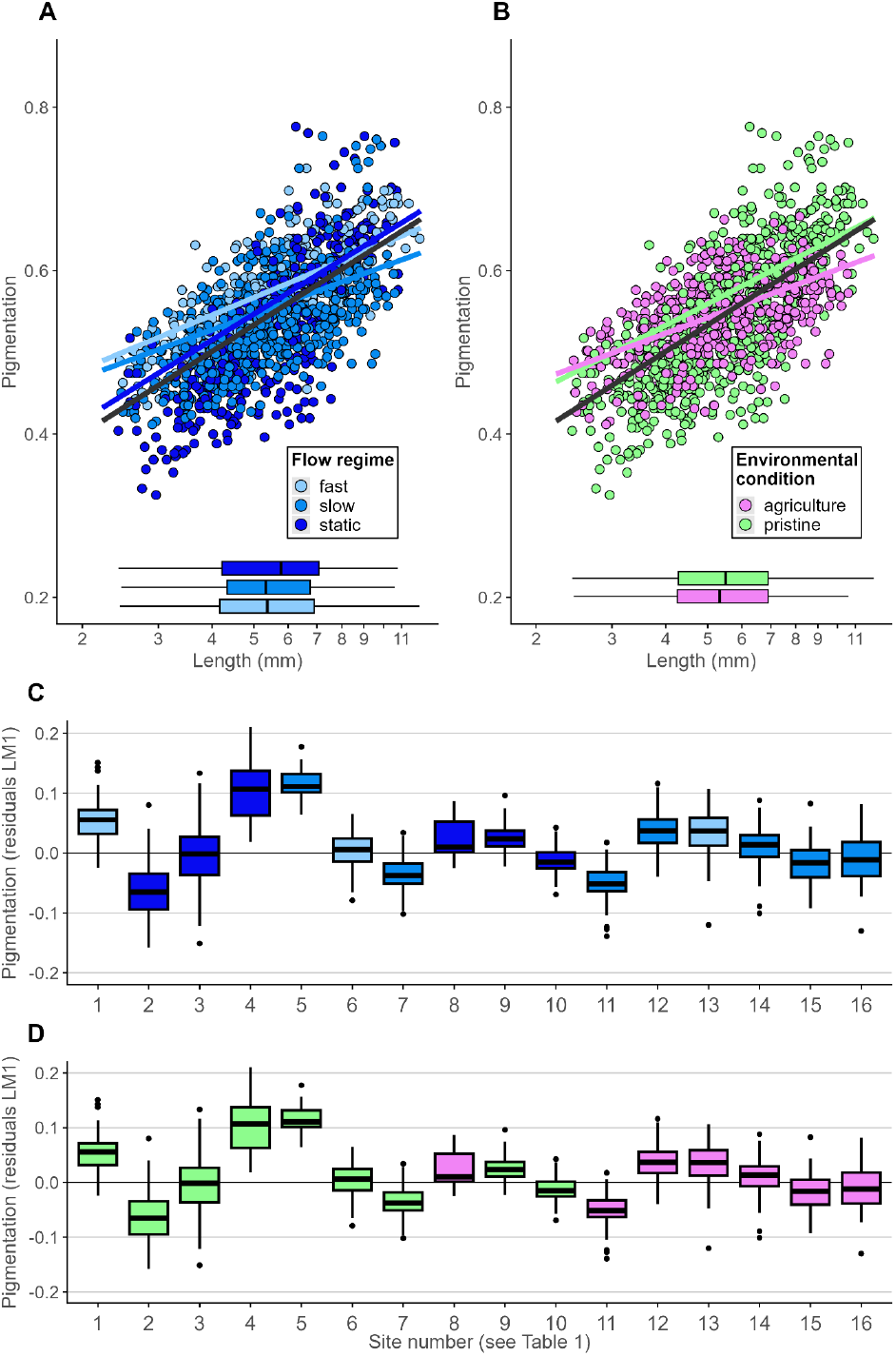
The relationship of body size and pigmentation grouped by flow regime and condition. We found interactive effects between body size and flow regime (A) and environmental condition (B) on pigmentation (see Table 2 for model results). Similar to habitat type (LMM 1), these models explain much less variation in the relationship of body size and pigmentation than the series of Generalized Additive Models (GAM, Fig 3, Table3) with continuous variables of substrate darkness and nutrition. Moreover, the effect is not consistent among sites belong to the same groups (C/D)

## Tables

**Table S1.** Amino acid concentration for all sites. Amino acid (AA) content of all sampled sites. The sum of all AAs (i.e., total protein) are reported as concentration in substrate mass (µg/g). Individual AAs are reported as proportion in total protein. The green color coding is row-based, with darker green indicating higher proportions and brighter green indicating smaller proportions.

| Site number | Site name | Habitat type | Sum (total protein µg/g) | Ala | Cys | Gly | Val | Leu | Ile | Nle | Thr | Ser | Asp | Pro | Met | Glu | Phe | Lys | Tyr | His | Trp |
| --- | --- | --- | --- | --- | --- | --- | --- | --- | --- | --- | --- | --- | --- | --- | --- | --- | --- | --- | --- | --- | --- |
| 1 | Asahikawa 03 | Forest runlet | 0.059 | 0.043 | 0.088 | 0.082 | 0.057 | 0.099 | 0.036 | 0.009 | 0.044 | 0.054 | 0.098 | 0.058 | 0.136 | 0.015 | 0.051 | 0.076 | 0.025 | 0.025 | 0.002 |
| 2 | Atsushinai River | Forest runlet | 0.043 | 0.039 | 0.147 | 0.075 | 0.054 | 0.088 | 0.038 | 0.012 | 0.042 | 0.059 | 0.102 | 0.058 | 0.131 | 0.005 | 0.048 | 0.076 | 0.017 | 0.008 | <0.001 |
| 3 | Kamekawa River | Forest runlet | 0.035 | 0.043 | 0.077 | 0.089 | 0.056 | 0.09 | 0.04 | 0.019 | 0.037 | 0.054 | 0.095 | 0.06 | 0.127 | 0.004 | 0.05 | 0.103 | 0.021 | 0.007 | 0.027 |
| 4 | Asahikawa 02 | Pasture runlet | 0.042 | 0.043 | 0.082 | 0.089 | 0.059 | 0.097 | 0.048 | 0.01 | 0.028 | 0.044 | 0.106 | 0.066 | 0.151 | 0.015 | 0.057 | 0.065 | 0.024 | 0.012 | 0.003 |
| 5 | Asahikawa 07 | Pasture runlet | 0.038 | 0.044 | 0.074 | 0.082 | 0.057 | 0.096 | 0.038 | 0.013 | 0.042 | 0.059 | 0.101 | 0.058 | 0.14 | 0.016 | 0.052 | 0.078 | 0.026 | 0.025 | <0.001 |
| 6 | Asahikawa 09 | Pasture runlet | 0.021 | 0.042 | 0.109 | 0.083 | 0.055 | 0.092 | 0.037 | 0.028 | 0.033 | 0.058 | 0.105 | 0.062 | 0.135 | 0.016 | 0.046 | 0.058 | 0.019 | 0.014 | 0.009 |
| 7 | Hiraoka Park | Pasture runlet | 0.033 | 0.049 | 0.1 | 0.092 | 0.056 | 0.083 | 0.035 | 0.016 | 0.03 | 0.064 | 0.106 | 0.064 | 0.141 | 0.009 | 0.042 | 0.08 | 0.017 | 0.015 | <0.001 |
| 8 | Teshio 03 | Pasture runlet | 0.026 | 0.045 | 0.088 | 0.083 | 0.055 | 0.09 | 0.036 | 0.019 | 0.053 | 0.065 | 0.101 | 0.057 | 0.129 | 0.005 | 0.043 | 0.096 | 0.014 | 0.021 | <0.001 |
| 9 | Asahikawa 01 | Roadside ditch | 0.017 | 0.034 | 0.163 | 0.07 | 0.045 | 0.077 | 0.03 | 0.029 | 0.034 | 0.046 | 0.082 | 0.057 | 0.114 | 0.017 | 0.042 | 0.056 | 0.018 | 0.012 | 0.074 |
| 10 | Asahikawa 04 | Roadside ditch | 0.038 | 0.038 | 0.216 | 0.066 | 0.044 | 0.086 | 0.029 | 0.028 | 0.032 | 0.053 | 0.087 | 0.049 | 0.123 | 0.013 | 0.045 | 0.042 | 0.021 | 0.01 | 0.019 |
| 11 | Nakahoronuka River | Roadside ditch | 0.041 | 0.045 | 0.101 | 0.081 | 0.05 | 0.085 | 0.031 | 0.012 | 0.039 | 0.05 | 0.115 | 0.055 | 0.142 | 0.016 | 0.047 | 0.082 | 0.021 | 0.025 | <0.001 |
| 12 | Asahikawa 05 | Concrete channel | 0.079 | 0.046 | 0.088 | 0.08 | 0.048 | 0.092 | 0.035 | 0.012 | 0.044 | 0.054 | 0.099 | 0.051 | 0.143 | 0.011 | 0.05 | 0.083 | 0.029 | 0.032 | 0.001 |
| 13 | Asahikawa 06 | Concrete channel | 0.05 | 0.048 | 0.103 | 0.091 | 0.054 | 0.089 | 0.035 | 0.012 | 0.019 | 0.033 | 0.14 | 0.054 | 0.171 | 0.012 | 0.051 | 0.063 | 0.019 | 0.006 | <0.001 |
| 14 | Asahikawa 08 | Concrete channel | 0.038 | 0.045 | 0.121 | 0.077 | 0.047 | 0.094 | 0.029 | 0.021 | 0.048 | 0.051 | 0.118 | 0.046 | 0.132 | 0.019 | 0.043 | 0.065 | 0.023 | 0.021 | <0.001 |
| 15 | Teshio 01 | Concrete channel | 0.065 | 0.044 | 0.139 | 0.076 | 0.048 | 0.06 | 0.03 | 0.009 | 0.047 | 0.057 | 0.094 | 0.048 | 0.148 | 0.01 | 0.049 | 0.091 | 0.024 | 0.024 | 0.002 |
| 16 | Teshio 02 | Concrete channel | 0.036 | 0.047 | 0.099 | 0.087 | 0.053 | 0.08 | 0.032 | 0.015 | 0.05 | 0.061 | 0.079 | 0.058 | 0.132 | 0.011 | 0.043 | 0.103 | 0.021 | 0.028 | 0.002 |

